# In vitro design of intrathecal drug administration therapies

**DOI:** 10.1101/2025.11.25.690590

**Authors:** Ayankola O. Ayansiji, Caleb Gardner, Sebastien Dors, Daniel S. Gehrke, Francisco Moral-Pulido, Konstantin Slavin, Andreas A. Linninger

**Affiliations:** University of Illinois at Chicago, USA; Visiting Research Student at University of Illinois at Chicago, USA; University of Illinois at Chicago USA

## Abstract

Due to the scarcity of reliable in vivo data, the pharmacokinetics of intrathecally (IT) administered drugs remain inadequately quantified. Designing new therapies is further hindered by variability in experimental methods, inter-individual and inter-species differences, and poor reproducibility across animal and human studies.

To address these limitations, we developed an anatomically accurate, subject-specific replica of the cerebrospinal fluid-filled spaces of the human central nervous system (CNS) using a multistep mold/casting process. The 3D-printed, transparent, deformable CNS phantom enables precise control of infusion and physiological parameters, allowing systematic generation of reliable and repeatable biodispersion data for lumbar IT infusion protocols. Pulsatile artificial cerebrospinal fluid (CSF) flow within the closed system was tuned to replicate subject-specific stroke volumes and flow rates observed in MRI. The model’s optical clarity facilitated high-speed visualization and tracking of tracer dispersion, exceeding the temporal resolution of current neuroimaging techniques.

Experimental series spanning physiologically relevant CSF and infusion conditions enabled quantification of the spatiotemporal distribution of IT-administered tracers. Inversion of the parabolic diffusion equation provided estimates of the coefficient of *effective dispersion*. A distributed pharmacokinetic model was used to evaluate the influence of chemical kinetics and mass transfer on tracer behavior. The proposed experimental apparatus for in vitro design of IT therapies offers a complementary or alternative approach to traditional trial-and-error animal studies.

## 2 Introduction

The blood brain barrier poses a formidable obstacle for drug delivery^1,2^. Intrathecal (IT) drug delivery is a minimally invasive modality which can overcomes this limitation^3–6^. Two administration modes are common: (i) chronic dosing with an implanted drug pump or (ii) acute infusion through a lumbar catheter. Drug pumps operating with slow, pulsed or steady infusion enjoy a long successful application history for chronic pain and spasticity^7–11^. As far back as 1980s, IT drug system has been in use for various indications such as spinal stenosis, discogenic pain, spasticity^7,9–12^. Acute administration, usually with short intervals at relatively high injection rates (1-3 ml/min), is a promising avenue for new therapies like antisense oligonucleotides (ASO) or enzyme replacement therapy targeting the brain^13–16^.

A major hurdle to the effective administration of therapeutics to cerebral and spinal targets via IT stems from the lack of data that correlate diverse infusion protocols and CSF dynamics to speed and localization of the active drug molecule. Quantitative methods to predict the effect of injection volume, infusate dose/dilution and injection speed are incomplete. Trial-and-error experimentation in humans to ascertain optimal infusion parameter is not feasible for several reasons. IT is invasive and associated with pain and infection risks even in the hands of expert clinicians^17,18^. It is almost impossible to carry out trials that systematically vary key parameters including (i) size and configuration of anatomical spaces, (ii) amplitude and frequency of CSF dynamics or (iii) administer multiple drugs with different molecular properties. Ethically, it is challenging to justify explorative trials in human subjects, yet parametric experimentation with repeats covering the extremes of physiological ranges is indispensable for robust statistical analysis.

To overcome these limitations due to data scarcity, we have created an in vitro model of the human central nervous system that replicates the main physiological and geometric characteristics of pulsatile CSF dynamics in humans. Our study centers on quantifying physical transport of IT drug administration subject to different injection modes and physiological parameter ranges of subject-specific CSF volume, pulsatile amplitude and frequency.

Data from systematic trials with multiple repeats enabled the quantification of CSF-mediated drug transport as a function of injection modes and natural CSF parameters. We precisely characterized the *effective* bio-dispersion coefficient as a function of CSF amplitude and frequency. Results suggest that tracer bio-dispersion matches the spatiotemporal dynamics of a *diffusive* dispersion process.

We conducted experiments with inert tracers to focus only on physical transport, avoiding interference from biological factors like uptake or decay. Chemical kinetics for specific drugs was considered a separate step, which we addressed using pharmacokinetic modeling. Integration of drug transport data with biochemical reactions will be demonstrated via a computational study at the end of this paper.

## 3 Methods

### 3.1. Manufacturing of a Functional CNS Model

An anatomically faithful, fully functional replica of the CNS was manufactured through a two-stage cast and mold process. We chose this technique over direct 3D printing, because casting allows for wide material choices including soft and transparent polymers that would not be directly printable. Deformability is a critical requirement to allow for natural CSF flow patterns inside closed anatomical spaces which is impossible to achieve in open rigid models. The five steps of the mold/casting process are presented in brief; a more detailed description is presented in Appendix A. A flowchart of the manufacturing stages and processes can be found in appendix B.

**Imaging and segmentation**. We acquired subject-specific anatomical properties of the human cranial and spinal SAS in several imaging sessions (see detailed MR protocol elsewhere^19,20^). The acquisition protocol of the human MRI data is presented in appendix A1; the original imaging data can be found in previous study^21^. DICOM image stacks of MR image data^20,22,23^ were segmented using ITK-SNAP^24,25,26^. The segmented data was structured into an inner and outer surface mesh encompassing the spinal SAS and stored as in stereolithographic (STL) format (with 53,514 elements) with good quality to retain anatomical details of the extracted images, especially paired bundles of nerve roots. The image reconstruction is detailed in appendix A2.

**Mold design**. We choose to design section molds in eight pieces to cover the entire neuraxis as shown in Fig.1. Each piece was designed to measure under 10 cm in length to fit into the regular commercial printing rig (Comgrow Creality Ender-5 3D Printer^27^). Ring-like grooves were added to the connecting interfaces of each section using Solidworks. The groove/connections facilitated precise spacing and tight connections required in the subsequent assembly. The section meshes in GCODE format were loaded into a 3D printer (Comgrow Creality Ender-5 3D Printer^27^). The total time for the printing of all segments was about 29 hours. Detail description of the mesh slicing process is shown in appendix A3.

**Fig. 1.**
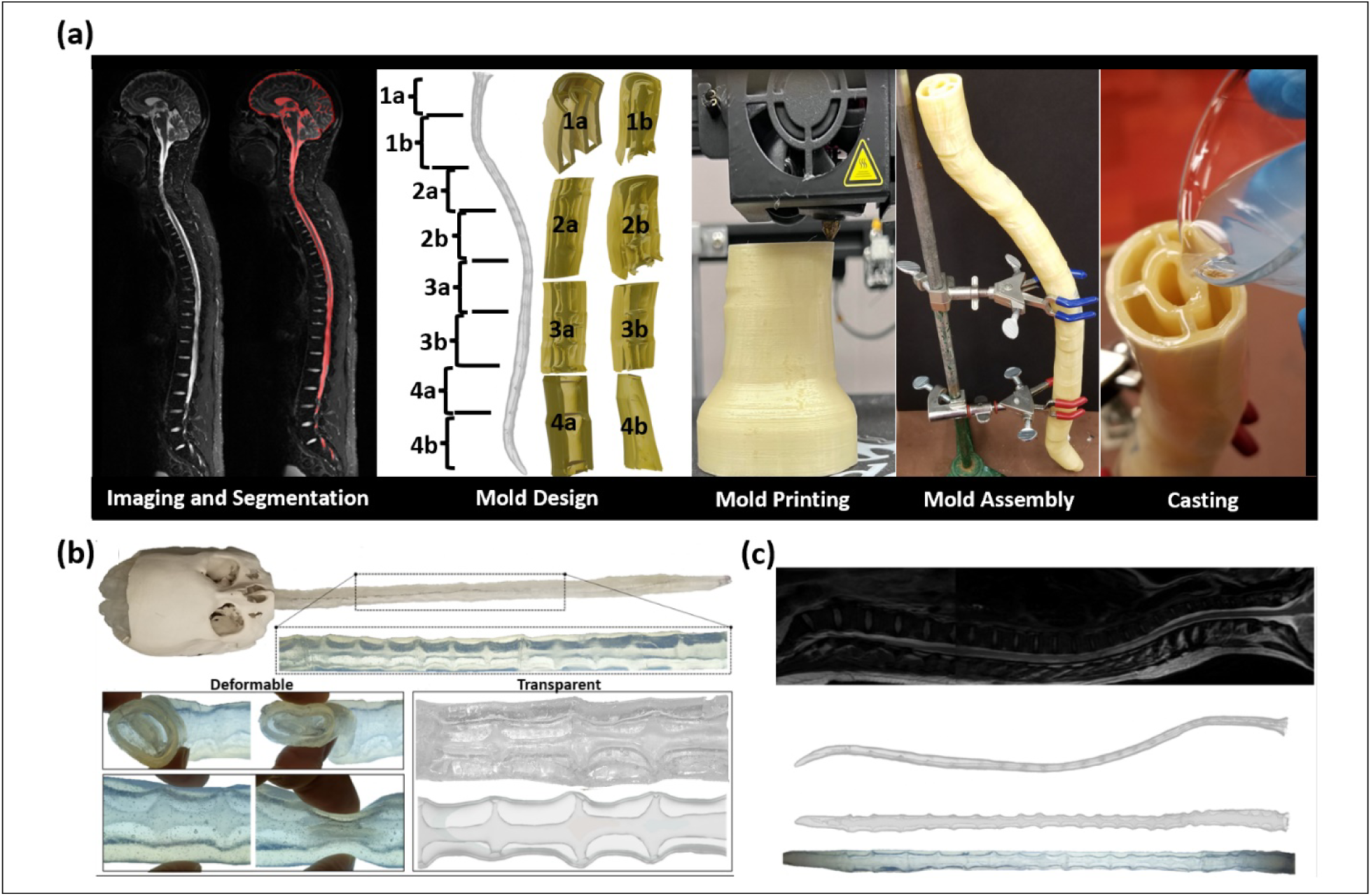
(a) The schematic diagram showing the five steps involved in the mold/casting process (i.e. Imaging and segmentation, mold design, mold printing, mold assembly, and casting). (b) The casting result shows the deformable ability of the cast spine and transparency of the spine which allows for proper tracking of tracer in the phantom during experimental activities. (c) The close reproduction of MRI data of the spine and the manufactured spine with the presence of anatomical features. The final phantom has dimensions very close to that obtained in MRI with average percentage difference in diameter and areas of 1.88% and 3.78% in the area.

**Mold Printing**. Each section in GCODE format was 3D printed with dissolvable polyvinyl alcohol (PVA) as filament molds using filament printing mode. PVA properties are listed in table 1 with detailed explanation of the printing including the printer settings are shown in appendix A4. The proposed sectioned design had the advantage of providing access to interior surfaces for smoothing and curating rough edges of raw molds. Rough sections were smoothened with a brush and water which ensured high surface quality of the mold as shown in Fig. A2 in appendix A, and final phantom.

**Table 1:** Table showing the characteristic properties of the PVA filament.

| Properties | Value/Dimension |
| --- | --- |
| Roll diameter | 3 (mm) |
| Filament Roundness | +/- 0.07 (mm) |
| Filament Diameter | +/- 0.05 (mm) |
| Net weight | 0.5 (kg) |
| Recommended Print Temperature | 160 - 190 ($^{\circ}$C) |
| Solubility | Highly soluble in water |

**Mold Assembly**. The eight section pieces were assembled and fastened with clear Elmer’s Glue (Elmer’s Products, Inc.) into a complete 3D mold of the spinal SAS (negative space). The alignment of the inner and outer mold pieces was precisely safeguarded by the printed spacer grooves to avoid breakage of the mold and to make it watertight for the casting process and left to dry for 24 hours (see Appendix A5).

**Casting**. TAP Platinum Silicone casting resin of shore A8 hardness (with Young Modulus of 0.75 MPa) obtained from TAP Plastics^28^ was poured in a way that 50mL of the resin enters the mold in 1 min (i.e. approximately 50mL/min flow rate) into the closed mold and allowed to settle. TAP is transparent after casting and endowed with the final CNS analogue (positive space) with the soft deformable tissue-like properties. The TAP has two sides, A (the base) and B (the catalyst), which are vigorously mixed and then poured into the assembled mold. The cast was left for 24 hours to ensure that it was fully cured. This material was chosen because it led to an elastic wave speed of 27.2 m/s, which is in the physiological range (3.5 – 33.8 m/s)^29–31^. More details on the casting process are given in detail in appendix A6. After curation, the entire mold/cast assembly was submerged in a water bath to gently detach and gradually dissolved the mold (negative) from the cast (positive space) by dissolving the water soluble TPA. Fig. 1c shows the manufactured phantom has the geometry of the real human spine as shown by the MRI data. Detail description of the mold dissolution is shown in appendix A7.

### 3.2 Functional dynamics reproduce natural CSF oscillations and enable tracer injection

**Pulsating Flow –**Pulsatile CSF motion in the human CNS is caused by oscillatory expansions of the cerebral blood compartment (possibly spread via the parenchyma) inside the rigid cranial compartment^32^; the resulting volumetric strain is transmitted from cranial to spinal CSF, which in turn expands the spinal dural spaces. Distensibility is an essential feature of our closed deformable phantom, which cannot be achieved in open rigid models. Accordingly, dural and pial boundaries of the CSF filled spaces are deformable and closed as shown in Fig. 1b. To induce pulsatile CSF motion, a piston pump in the head compartment was used to expand and contract a water filled ballon mimicking a distensible blood/brain compartment. The ballon’s pulsations emulate the expansion of vasculature which in turn sets in motion CSF flow from the cranial SAS into the elastically deformable spinal SAS.

In vivo, the volumetric expansion of the closed spinal SAS necessarily induces graded volumetric flow rates in the spinal CSF with snapshots in systole and diastole shown in Fig 2b. Because rigid models require open sacral and cervical sections, they enforce constant CSF flow along the neuraxis. The absence of deformability prevents graded flow amplitudes, resulting in unrealistic dispersion patterns. The proposed functionality enables us to cover the physiological range of stoke volume from 0.5-1 ml/beat^3,33^. The dynamic area deformation and the pressure signal that drives the dispersion process are plotted in Fig. 2c (obtained by simulation). The average change in area is 0.248mm^2^, the average radial deformation is 1383µm, average flow change is 0.78ml/sec, and the average change in pressure is 2.72 Pa. Pressure signals were measured in vitro using water manometrics with plots in Fig 2d.

**Fig. 2.**
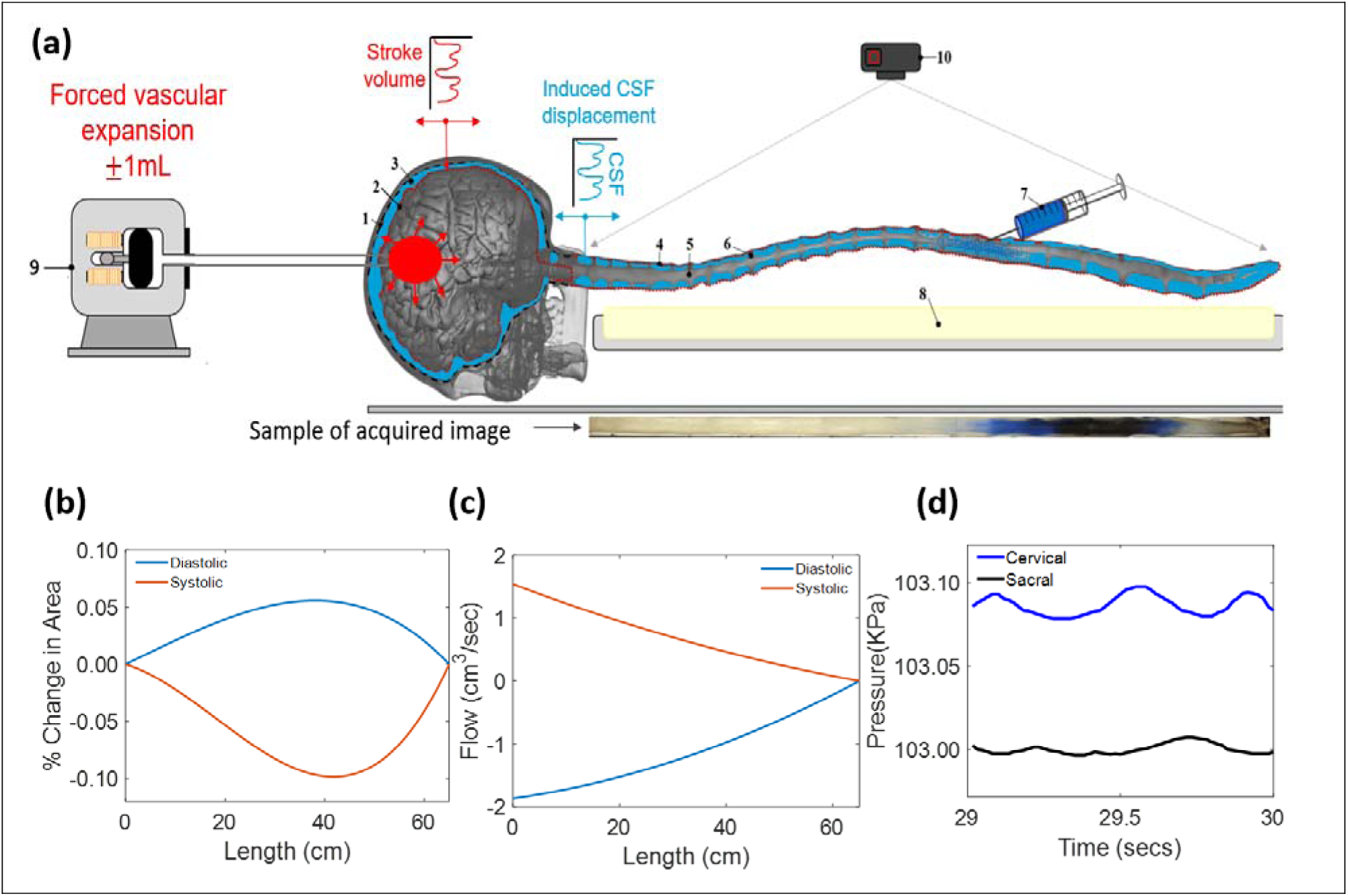
(a) Diagram of the CNS spine model in position on the benchtop for testing of infused tracer bio dispersion. Where part (1) Rigid Cranial Vault (2) Deformable Brain Parenchyma (3) Idealized Cerebrospinal Fluid (4) Deformable Dura Matter (Subject Specific (S.s)) (5) Spinal Cord Bundle (S.s) (6) Peripheral Nerve Bundles (S.s) (7) Infusion Source (8) Illumination Source (9) Piston Pump (Oscillations) (10) Optical Recording Camera. (b) Simulated area, (c) Simulated flow. The flow is attenuated along the length of the neuraxis. (d) measured pressure at different times. The temporal variation of the cross-sectional areas shows small canal deformations, which allows the pulsatile flow inside the distally closed CNS phantom. In addition, the closed model induces attenuated flow amplitude from the cervical towards the sacral region as shown in (c) for diastole and systole.

**Tracer infusion system–** For acute tracer infusion, we used as programmable syringe in Fig. 2b to inject desired injection volumes at variable infusion rates. In the experiment, we injected 2ml of tracer for a period of 1 minute. Of note, our injection system is identical to clinical needle infusion requiring no valves, because the dura in the phantom is soft, water-tight and self-sealing.

**Optical tracking of the tracer fronts –**After injection for 1 minute, the syringe injection pump is stopped; further tracer motion occurs only due to natural CSF pulsation. Translucency of the dural surfaces allowed optical tracking with enough temporal resolution (120 fps) of intensity curves as a function of position along the neuraxis and time, see Fig. 4a.

**Analysis of dispersion data**. Videos files showing tracer intensity plots were processed with video analysis software^34^. In-house build MATLAB code was used for reconstructing the tracer intensity from the snapshots at different times as shown in Fig. 4b. Our experiments did not allow accurate determination of radial intensity gradients, which appeared to be small. Therefore, radially averaged intensities^21^ were plotted. Because of asymmetry in biodispersion due to lumbar injection and unequal size of the sacral and the cervical SAS, tracer profiles are not symmetric even in a perfect diffusion process. To account for skewed distributions, we designed a rigorous parameter estimation process termed *inversion of moments*, as an improvement over the method of moments (MoM)^21^ We defined the *effective* biodispersion coefficient as the diffusivity of an ideal diffusion process that best matches the time course of the asymmetric second moment, Left-sided second moment, *M*_2,*Left*_(*t*).

**Inversion of Moments.** Due to asymmetry and boundedness of the spinal CSF spaces, second moment trajectories progressively taper off when the concentration profiles reach the sacral region. To overcome this departure from ideal diffusion (i.e. symmetric, infinite domain), we dynamically invert the time dependent diffusion equation^35^ in eq (2) to obtain an effective dispersion coefficient, *D*, which optimally matches the evolution of simulated moments to the non-linear trends observed in anatomical spaces. The inversion process is able to handle curved moment plots, thus giving more accurate estimates of the dispersion coefficient than MoM. We chose moments in the objective of eq (1) instead of raw concentration profiles to suppress uncertainties from noisy tracer intensity acquisitions. The proposed moment inversion process (MIP) has thus integrative smoothing characteristics.

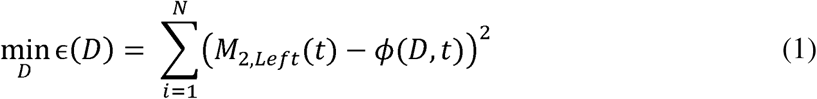

The ɛ(*D*) is the minimized error as a function of simulated diffusion, φ(*D,t*) is the left-sided second moment of the equivalent diffusion process for a given diffusion coefficient, *D*, at time dispersion coefficient, *D* in (1), the simulated moments φ(*D,t*) need to be computed by solving point, t., and *N* is the number of time period considered. In order to obtain the desired optimum the parabolic diffusion equation (2) which is achieved numerically using finite volume method^36^ as shown in appendix C3.3. The optimum value that minimizes eq (1) is taken to be the estimated *effective* dispersion coefficient. The non-linear inversion problem in Eq (1) was solved with MATLAB *fminunc*.

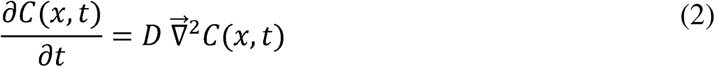

Here *C* is the concentration, *t* is the time, *D* is the *effective* dispersion coefficient. Experimentally observed left moments *M*_2,*Left*_(*t*) in (1) are calculated using experimental image data using Eqs (3-4).

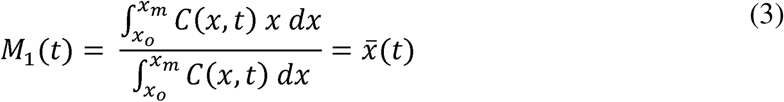

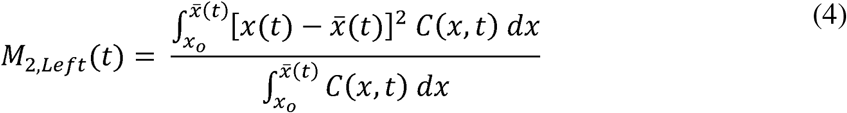

Also *x* is the position along the neuroaxis; *x*_O_ and *x*_m_, mark the lower and upper limit of the region of analysis. In each time frame, t, the first moment, *M*_1_(*t*) gives the location of the center of gravity, *x*(*t*); *M*_2,*Left*_(*t*) is the left sided mean spread of the visible concentration profile around its center, *x*(*t*), obtained in the experiment. The results in Appendix C3.2 table C1 show that inversion (MIP) was more accurate than the method of moments (MoM). This moment is *x*(*t*) i.e., where *x ≤ x*(*t*). The least square error as a function of dispersion coefficient, ɛ(*D*) determined by integrating over intensities to the left (in cranial direction) of the mean position, between observed left-sided second moment *M*_2,*Left*_(*t*) and the left-sided second moments of a simulated *equivalent* diffusion process (φ*_i_*) was minimized. The value of diffusivity in the model that attains the least square difference between observed and predicted moments as the objective in Eq. (2). Eq. (3) and Eq. (4) gives the left-sided second moments of the concentration profile.

In addition, the first moment, *M*_1_(*t*) of intensity plots along the neuraxis was computed as a motion and its speed [cm/min] was inferred from the slope of the *M*_1_(*t*) curve over time which function of time (total of 10 minutes at 1 minute interval) with the use of Eq (3). Caudocranial were obtained from the acquired intensity curves.

## 4 Results

### 4.1. Geometry validation of mold and cast

To validate the geometric fidelity of the manufacturing process, Table 2 shows the cross-sectional geometry comparison between the MRI data, printed mold and the final phantom. Object dimensions were obtained using the measurement tool in 3D Builder and verified with a physical ruler. The final phantom has dimensions very close to that obtained in MRI with average percentage difference in diameter and areas of 1.88% and 3.78% in the area, respectively. Fig. 1b qualitatively shows that the phantom matched anatomical features of the subject-specific MRI data. Gross geometrical dimensions of the dura, nerve roots and S-like ondulation of the CSF spaces along the neuraxis agreed with the anatomical features of the real human spine. The relatively small distortions were realized thanks to flow rate control (slow pouring at 50mL/min) of the casting material into the PVA mold.

**Table 2:** Table validating the manufactured spine dimensions at different locations (CM cisterna magna, T2 thoracis, L1 Lumbar, and S3 Sacral).

| Position | CM = 0 cm | T2 = 20 cm | L1 = 50 cm | S3 = 65 cm |
| --- | --- | --- | --- | --- |
| In vivo diameter (cm) | 1.30 | 0.70 | 1.00 | 0.78 |
| Mold diameter (cm) | 1.32 | 0.71 | 1.02 | 0.80 |
| Phantom diameter (cm) | 1.32 | 0.71 | 1.02 | 0.80 |
| % Difference in diameter* | 1.54 | 1.43 | 2.00 | 2.56 |
| Change in diameter (mm)*** | 0.01 | 0.02 | 0.005 | 0.003 |
| In vivo area (cm<sup>2</sup>) | 4.90 | 1.43 | 2.51 | 1.47 |
| Mold area (cm<sup>2</sup>) | 5.01 | 1.49 | 2.63 | 1.53 |
| Phantom area (cm<sup>2</sup>) | 5.01 | 1.49 | 2.63 | 1.53 |
| % Difference in area** | 2.24 | 4.03 | 4.78 | 4.08 |
| Change in area (mm<sup>2</sup>)*** | 0.20 | 0.49 | 0.10 | 0.05 |

\*Obtained by comparing the in vivo diameter with that of the manufactured phantom.
\*\*Obtained by comparing the in vivo surface area with that of the manufactured phantom.
\*\*\*Obtained by simulation

#### 4.1.1 Dimensions of the Human Spine Cross-Sections

The geometry of some selected cross-sections of the human spine are shown in Fig. 3a and their dimensions (minor and major radii) are presented in Table D1 in appendix D. The sampled positions along the subarachnoid space are Cisterna Magna (CM), cervical region (C3, C5, and C7), thoracis region (T2, T4, T5, T7, T9, T11), lumbar region (L1, L2, L5), and the sacral region (S3) as shown in Fig. 3d with their respective positions. The cross-sections are approximately elliptical with b and a as the major and minor radii, respectively. Fig. 3c shows the percentage of the computed cross-section area that is occupied by spinal tissue. Table D1 in appendix D also shows the CSF occupied area. More views of the cross-sectional areas are presented in appendix F.

**Fig. 3.**
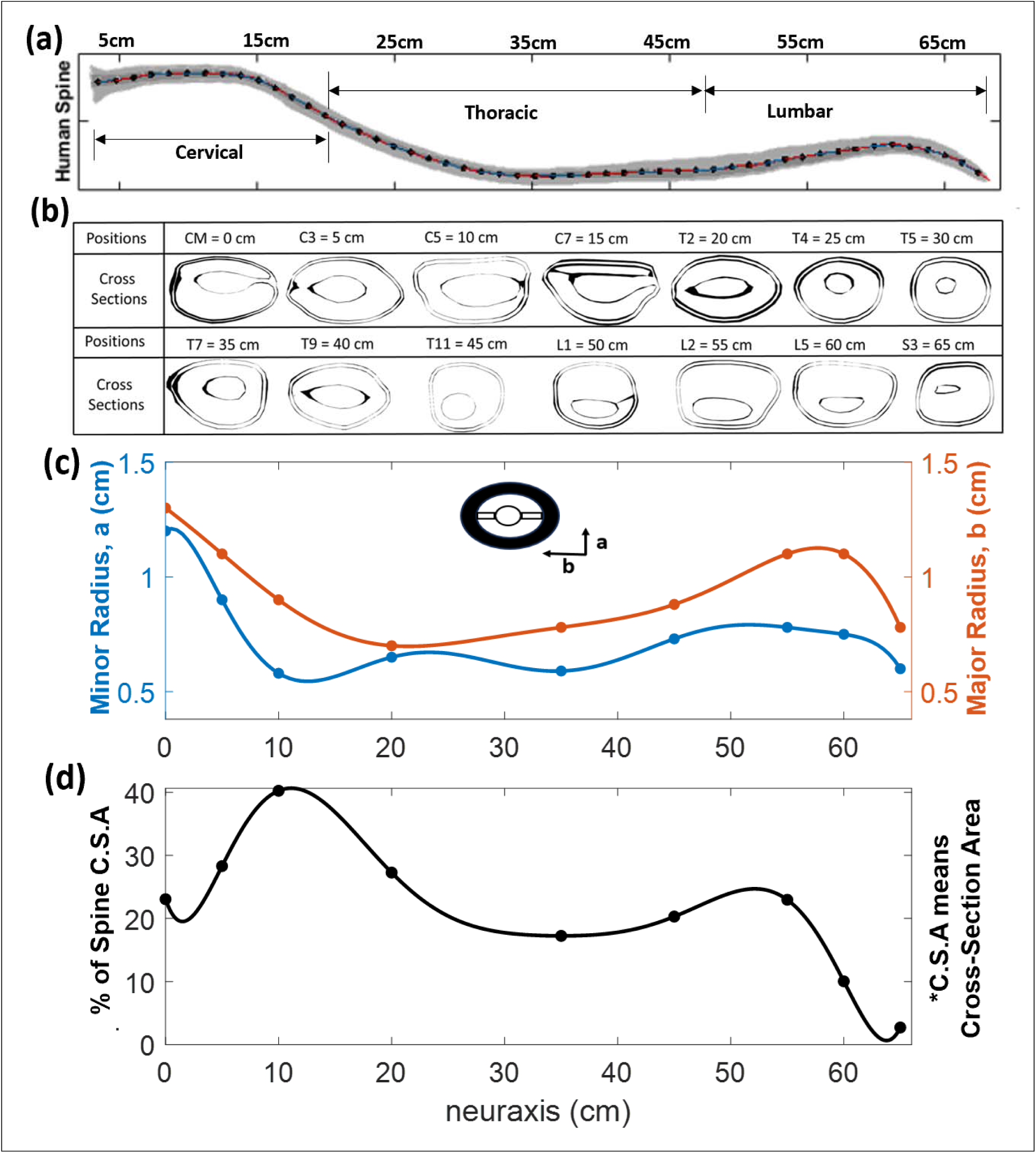
(a) The human spine with the regions. (b) The figure showing the different cross sections along the neuraxis at different positions. The ellipse shows the reference orientation for the major and minor radii. (c) The plots of the minor radius (dorsal to ventral extent) and the major radius (right to left extent of the CSF space) distribution along the neuraxis, respectively. The cross-sections whose data were used for the interpolation are also placed at the corresponding points. The formula for the major and minor radii is listed in appendix E. (d) The percentage of the cross-section area covered by the spinal tissue. This data is needed to assess the CSF covered area fraction in each cross section.

### 4.2. Anatomically Accurate Deformable CSF Model of the CNS

The closed deformable CNS analogue replicates pulsating CSF flow in the spinal canal. Transparency of all system boundaries enabling optical tracking of the tracer as shown in Fig. 2a. Cast specimens are durable (one phantom lasts about 100 experiment runs) adequately withstanding CSF flow oscillations without tearing or leakage. Average area deformation, pressure change, and flow change are 0.0021cm^2^, 0.0204 mmHg, and 0.78cm^3^/sec respectively (inferred by numerical simulation in terms of imposed stroke volume).

### 4.3. Systematic Study of Tracer Dispersion in IT Administration

#### 4.3.1 Effective Dispersion

We optically tracked the tracer intensity in IT infusion experiments (N=26) using trypan blue (from Millipore Sigma) using the setup in Fig. 2a. A wide parameter range matching physiological and infusion modes was covered: CSF frequency range was 40bpm to 127bpm. CSF cervical stroke volume was in the range of 0-1 mL/beat, most experiments were run at 0.5 or 1.0mL/Stroke. All experiments (N=26) were repeated three times; repeats typically deviated from prior runs less than 0.13 cm^2^/min.

Following the experimental set-up shown in Fig. 2a., 2 ml of the tracer (trypan blue) was injected into the spinal canal of the human spine phantom which is holding water or artificial CSF (aCSF) at room temperature. IT injections lasted 1 minute (phase 1) after which the syringe pump was stopped (phase 2). After stopping the injection, the tracer was allowed to be dispersed under the influence of the pulsation Fig 2a. Both phase-1 and phase-2 occurred in the presence of natural CSF pulsation.

A digital camera was used to record a video of the dispersion process (phase 2) over 10 minutes. In all our IT experiments, we observed that the tracer tends to move towards the cervical regions and the brain. The trypan blue intensity curves can be seen clearly expanding, initially almost symmetrically, from the lumbar injection site (x=0) in axial direction towards the sacral and thoracic region. Once the concentration front hits the sacral boundary, dispersion profiles become skewed and a caudocranial trend is clearly observed as evidence by a gradual shift of *M*_1_(*t*) from the injection site towards the cervical region. Snapshots taken at each minute as obtained from the recorded video are shown on Fig. 4a. The intensity profiles of the acquired frames are noisy so that direct inference of the dispersion coefficient is not advisable.

**Fig. 4.**
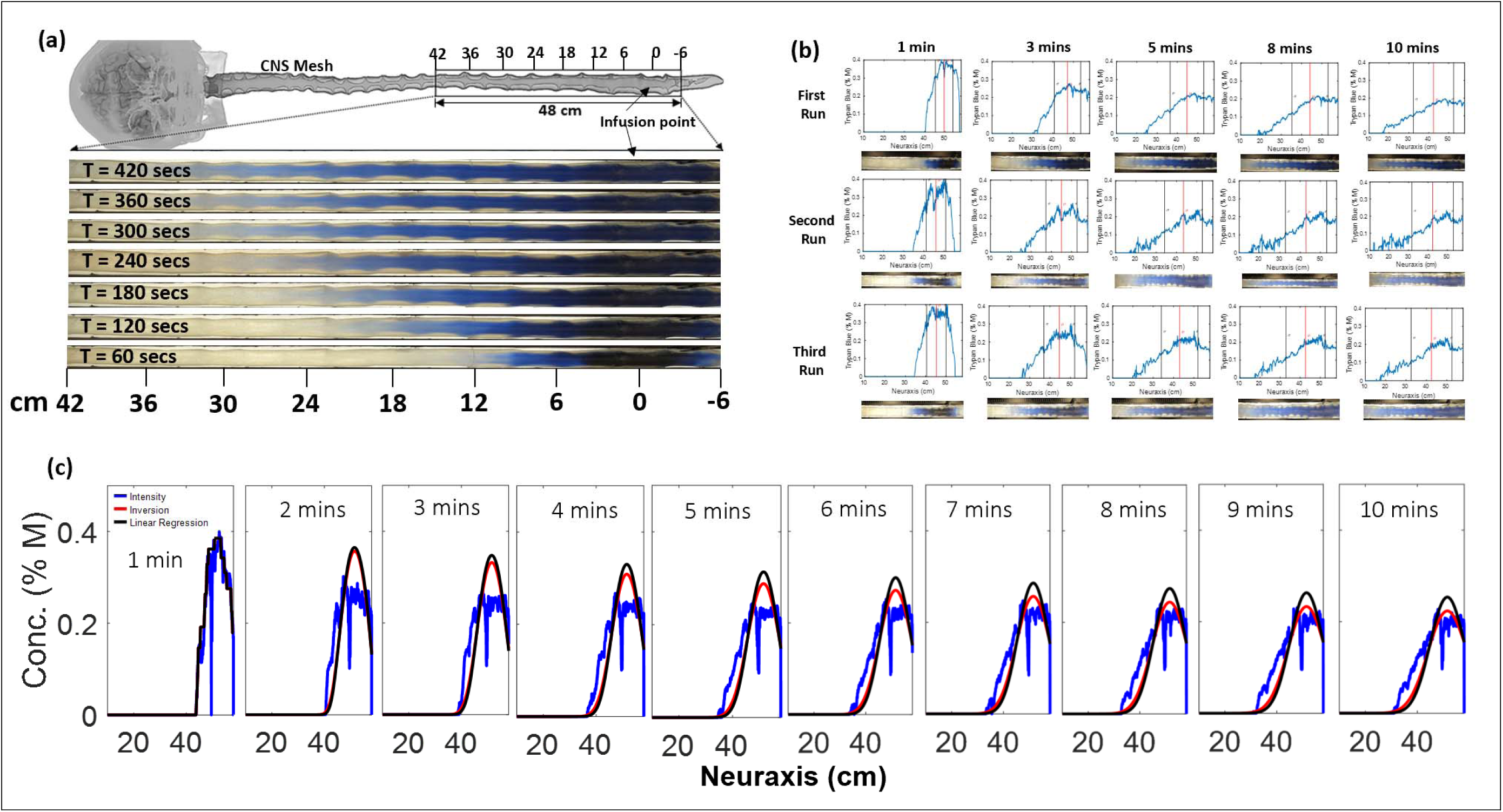
(a) Temporal evolution of the tracer dispersion (trypan blue) along the deformable model. Snapshots are taken each minute after the injection is already finished (t = 60 s). x-axis label indicate positions along the neuroaxis, point 0cm marks the infusion point. quotes of 42 cm (cervical) to −6 cm (sacral) (b) Typical raw concentration profiles obtained for different runs of the experiment at the frequency of 72 bpm and stroke volume of 0.5 mL/Stroke. (c). The plot shows the concentration profile predicted through the dispersion coefficient obtained by inversion method (shown in red), and method of moment (linear regression, in black). The blue color curve shows the raw intensity curves acquired from the experiment for trypan blue at frequency of 40bpm and 0.5mL/Stroke. Trends by inversion fit raw data better than those obtained by the method of moments.

The evolution of the first moment (the red vertical lines in Fig. 4b) representing the center of gravity of the curve is indicative of caudocranial motion of the tracer. The inversion process gave the dispersion coefficients with the use of least square method, see eq (1). Fig. 4c shows the plot of the second moment for different runs which shows that the second moment increases monotonically with time for the experiment at the frequency of 72bpm and stroke volume of 1.0mL/Stroke within the region of interest (ROI) of 10cm to 58cm. The observed dispersion coefficients for the three runs using the inversion method are range of D=6 – 8 cm^2^/min. Systemic parameter variation was performed for over 70 experiments. The dispersion coefficients obtained (for Trypan Blue) using the method explained in section 3.2 are shown in appendix G. Since the distribution of the tracer is solely based on the convective dispersion produced by the CSF dynamics, we expect the molecular diffusivity a thus the molecular weight to play a minimal role in dispersion. A theoretical basis for this assumption has been given by Moral-Pulido et al^37^.

Fig. 4c shows the compares intensities to predicted concentrations using MoM^21^ with results obtained through inversion (MIP). Although both methods exhibit similar overall trends, the concentration profiles using MIP values more closely match the acquired raw intensity data than those obtained using MoM.

The results in Fig. 5 summarize the dispersion coefficients as a function of CSF amplitude and frequency obtained in a set of experiments (N=26). The data also shows the reproducibility of the second moment trends using the inversion method.

**Fig. 5.**
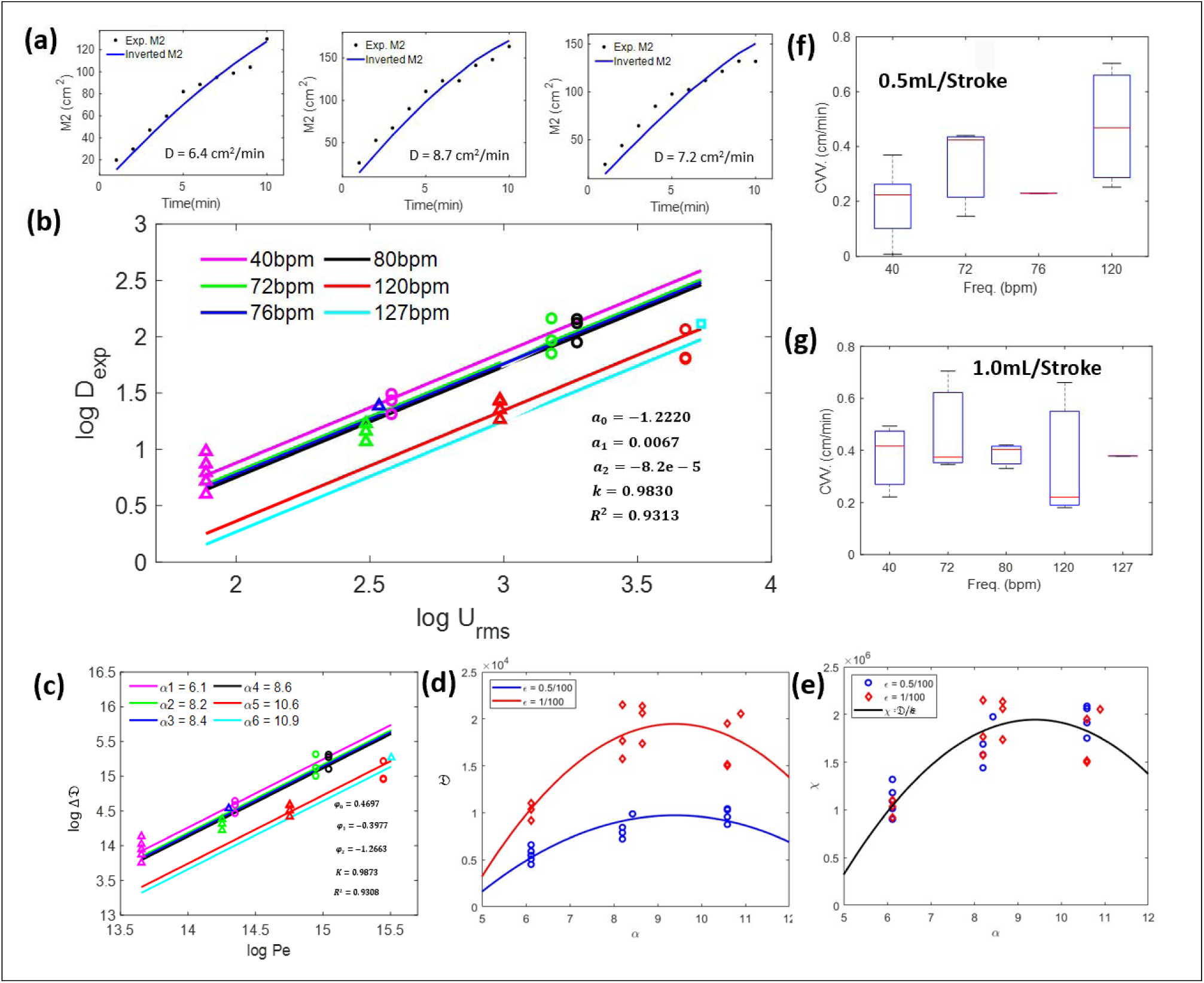
(a) Plots of second moments for the experiments and those from inversion process obtained in three repeated runs. The inverted second moment follows the trend of those obtained from the experiment (b) The relationship between the dispersion coefficient and frequency in dimensional and (c) dimensionless form for 26 experiment (d) Dimensionless dispersion coefficient prediction curves, as functions of the Womersley number (α) and the amplitude ratio (ε) (e) Normalized dimensionless dispersion coefficient prediction curve, as a function of the Womersley number (f-g) Caudocranial velocity (CCV) changes of the tracer relative to frequency for pulsation for 0.5 mL/beat (f) and 1.0 mL/beat (g).

Fig. 5a shows three repeated runs of the experimental second central moments compared to the predicted moments obtained using the inferred (i.e., optimal) effective dispersion for the experiment conducted at a frequency of 72bpm and a stroke volume of 1 mL per cycle. The simulated moments derived through inversion successfully capture the nonlinear, tapered trends observed in the data, which are attributed to anatomical asymmetries along the neuraxis affecting intrathecal (IT) tracer dispersion. The temporal evolution of experimental and predicted moments shows good agreement. Furthermore, all three experimental repeats under identical parameters (72bpm, 1 ml stroke volume produced consistent moment trends and inferred dispersion coefficients (D = 7.4 ± 1.9%). The similarity in both the experimental moment profiles and the estimated coefficients (D –D) highlights the reproducibility of the MIP-based method, despite unavoidable variability in lighting conditions between experiments.

**Functional dependence of the effective dispersion.** The plots of Fig 5b show in raw dimensional form a clear correlation between the speed of dispersion and CSF stroke volume (amplitude of CSF pulsations expressed in terms of root-mean-square velocity, *U_rms_*, as defined in Eq. H6 in Appendix H. The dispersion coefficients, *D*, increases with CSF amplitude. Note also that repeat experiments show data points with the same abscissa, which gives a measure the repeatability in the experimental procedure. Typical variance between repeat experiments was 2.13%. Parametric dependence of CSF frequency is also encoded as parallel lines.

The sequence of Fig. 5c-e summarizes the relationship between the *effective* dispersion coefficient in terms of CSF amplitude and frequency in dimensionless form. In dimensionless form, two correlations have been implemented, modeling the dispersion as a function of the non-dimensional numbers referring the frequency and the amplitude. The first correlation uses suitable logarithmic scales, showing that the logarithm of the effective dispersion is linearly dependent on amplitude with vertical shifts dependent on CSF frequency (ordinate offset). The second correlation uses scale analysis in the transport equation, which models the effective dispersion as a product of both amplitude and frequency effects of CSF motion.

**Experimental dependence on root-mean-square velocity and frequency.**

A statistical correlation of the listed experimental dispersion coefficient (in appendix G), D_exp_, as a function of natural CSF oscillations in terms of amplitude, root-mean-square velocity, *U_rms_* and frequency, *f*, on a double logarithmic scale is given in Eq. (5). A quadratic frequency dependence was also incorporated as an f-dependent offset, with 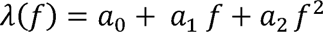 with *a*_0_ = −1.2220, *a*_1_ = 0.0067, and *a*_2_ = −8.2069 x 10^−5^ and drawn parametrically to Fig 5b. *K* has the value of 0.9830. Taking advantage of the logarithmic form of eq (8), all parameters could be estimated simultaneously by linear regression with coefficient of determination for the best fit of R^2^=0.9313.

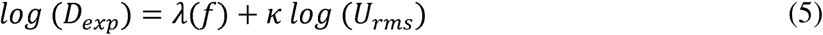

**Experimental data in terms of dimensionless numbers (Womersley and Peclet)**

Data could also be expressed in dimensionless form as shown in Eq. (6). The dimensionless effective dispersion changes, 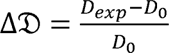, were plotted in Fig 5c based on raw Data could also be expressed in dimensionless form as shown in Eq. (6). The dependence of dispersion change, *log*(ΔD),, on the dimensionless oscillatory flow amplitude dimensional data of Fig 5b. The logarithmic correlations permit linear fitting for quantifying (Peclet number) and dimensionless frequency (Womersley number). Here, α is the Womersley number, 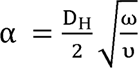 is the hydraulic diameter, 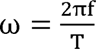 is the frequency in Hertz, *T* = 60 secs is the period, *D*_O_ = 1.938 x 10^−6^ cm^2^/min, and υ is the kinematic viscosity.

The choices of dimensionless characteristics follow prior approaches used to describe transport in oscillatory flows38.

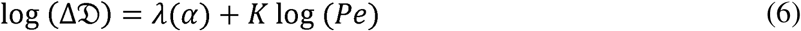

The best fit parameter set (K=0.9873, φ_O_ = 0.4697, φ_l_ = −0.3977, and φ_2_ = −1.2663, in Eq. (6) has a coefficient of determination of R^2^ =0.9308, where 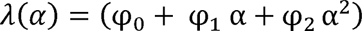 dispersion coefficient stabilizes above a critical value of *α*, followed by a slight reduction of this Fig 5c. In the case of the frequency, it is noticeable that the increment is not monotonous: the parameter.

**Scale analysis.** In addition to the statistical analysis of raw experimental data in dimensional form (in Eq 5), *scale analysis* of the advection-diffusion transport equation was performed to predict the *effective* dispersion coefficient (see details in Appendix H). Two key parameters emerged from the non-dimensional transport equation: (i) the Womersley number, *α*, and (ii) the pulsatile amplitude ratio, ɛ = (*U*_rms_/ω)/*L*. Scale analysis underscored the experimentally observed significance of amplitude and frequency, but introduces dimensionless volumetric deformation ratio, *ɛ*, thus pinpointing the value of the volumetric stain of the entire CSF filled space as a characteristic number. Thus, the *effective* dispersion coefficient is also parameterized as a function of these two characteristics in Eq (7),

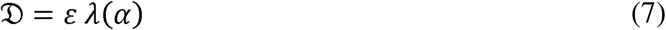

where 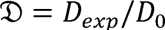 is the non-dimensional dispersion coefficient, and 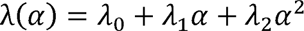 is the Womersley-dependent function, with λ_0_ = −5477 x 10^3^, λ_1_ = 1580 x 10^3^, and λ_2_ = −84.09 x 10^3^ as best fit. This fit yields a coefficient of determination

R^2^ = 0.9069, a match similar to that of the statistical model in (6). Additionally, the dispersion coefficient can be rewritten in a *normalized* form, 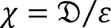, yielding a unique Womersley-dependent function. The dependence of the effective dispersion as function of both parameters is presented in Fig. 5d-e: both panels show an increment of the dispersion coefficient within the Womersley number, achieving its maxima at α = 10. However, the amplitude ratio scales the dispersion coefficient in an order-unity factor.

The comparison of the three models shows that the two main governing effects are related to the CSF pulsations, represented by its frequency (Womersley number, *α*), and its amplitude, The comparison of the three models shows that the two main governing effects are related to represented by its average velocity *U*_rms_ (or *P_e_* and *ɛ*). The dependence of the *effective* dispersion as a function of both parameters is presented in Fig. 5b-e: both approaches show the amplitude of CSF motion scales the phenomenon, modulated by the flow frequency effect.

Each function presented in Eq (5)-(7) can be used to predict *effective dispersion* along the neuroaxis as a function of subject specific anatomical and CSF motion parameters (e.g. pulsatile frequency and amplitude). Quantification of the physical dispersion speed based on this experimental series closes an important knowledge gap for the design of IT therapies. The integration of dispersion with biochemical kinetics will be shown in section 4.5.

#### 4.3.2 Speed of caudocranial motion

The caudocranial velocity (CCV) was also computed as the change in first moment with time. Eq. (7) and Eq. (8) shows the correlation between the CCV and the frequency, *f*, and root The caudocranial velocity (CCV) was also computed as the change in first moment with mean square velocity, *U*_!ms_, respectively.

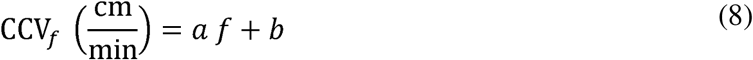

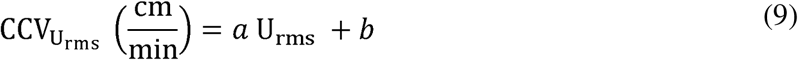

Where *a* and *b* are 0.0018 and 0.2144, respectively, for Eq. (8) and 0.0046 and 0.2650 respectively, for Eq. (9).

Fig. 5f and g show the relationship between caudocranial velocity and oscillating frequency for both stroke volumes. As seen, there is a remarkable increment in the CCV, especially for the minimal stroke volume. However, increasing the CSF amplitude reduces the frequency-dependent effect, becoming weak at higher frequencies.

### 4.4 The Effect of Nerve Roots on Tracer Dispersion

We conducted experiments to quantify the effect of nerve roots on the speed of tracer dispersion. For this purpose, we manufactured a second CSN model in which dentrite ligaments and nerve root bundles were absent as shown in Fig. 6b (spinal canal without nerve roots), giving rise to a quasi-annular spinal SAS. Fig. 6b shows the experimental results of dispersion of tracer in the spinal canal with inclusion or absence of microanatomical features. As evident in Fig. 6b, the tracer spreads faster in the spinal model with nerve root than that without nerve roots. Biodistribution was 321.1% faster when nerve roots were present. The experimental results confirm our earlier experimental findings^3^ as well as predictions obtained with direct numerical simulation^21,39–41^ where it has been reported that nerve root increases the dispersion of the tracer in the human spine due to the phenomenon called *geometry induced mixing*. Fig. 6c shows the quantitative representation of the effect of nerve roots. The 3-4 fold dispersion increase due to presence of nerve root accelerates drug dispersion under natural CSF oscillations. This benefit is active in after high-volume infusion as well as under slow infusion via a drug pump.

**Fig. 6.**
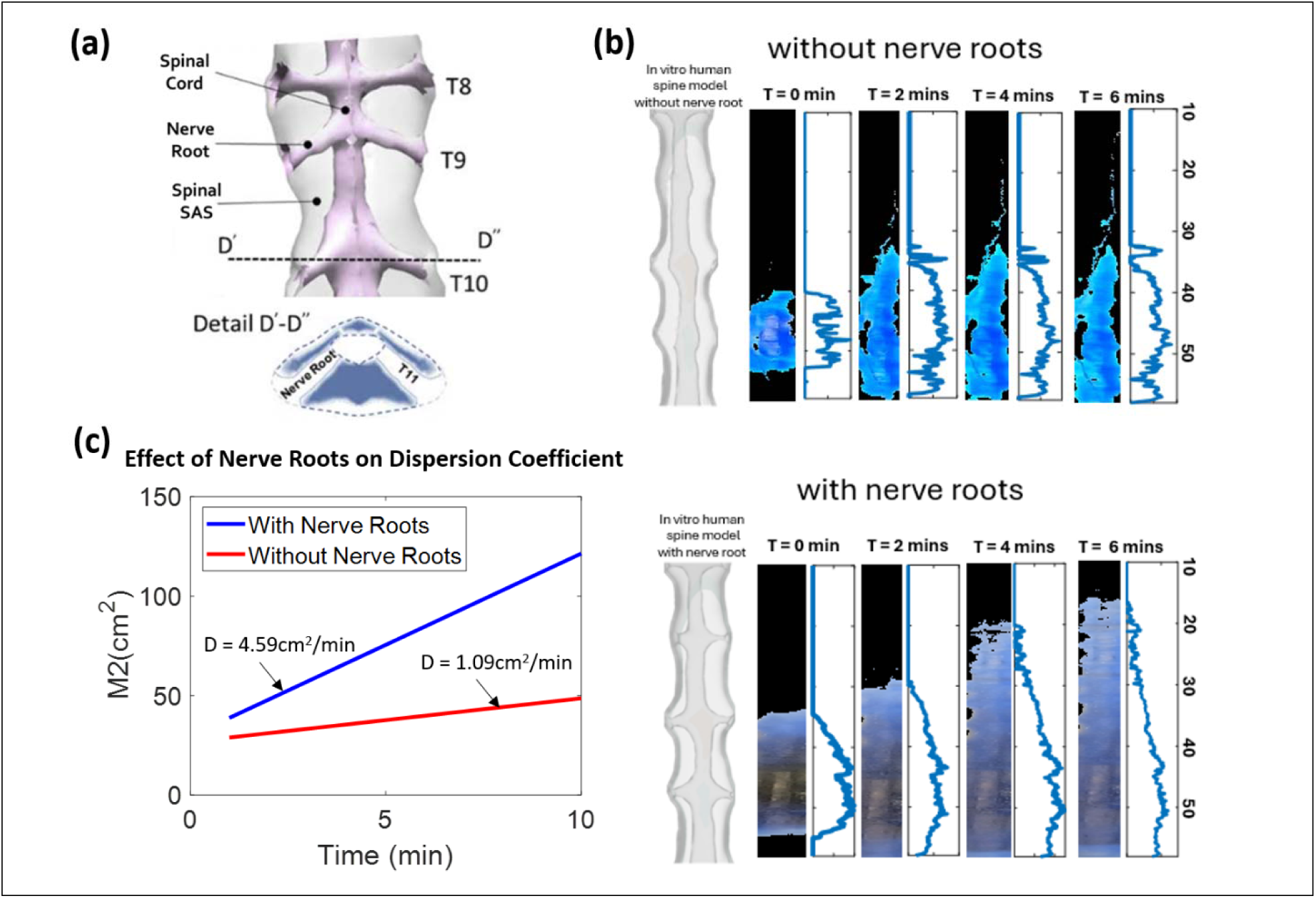
(a) Segment of the spinal SAS showing the detailed presence of the nerve root in the human spine. The line D’-D’’ shows the detail of the nerve root in the spine. (b) Effect of nerve root on the dispersion of the tracer. The system with nerve root has more than 4 times faster dispersion than that in which nerve roots were omitted in the manufacturing process. (c) Relation of the dispersion in a CNS spine model with peripheral nerve roots and without peripheral nerve roots.

### 4.5. Therapy design

How can physical dispersion data from this study be integrated with chemical kinetics and tissue uptake? We propose to predict drug biodistribution and uptake after IT using a distributed mechanistic pharmacokinetic model (PK). A schematic diagram of PK with six compartments (C1 – C6) introduced in a previous study^36^ is depicted in Fig. 7a. It enables simulation of injection and dispersion of active agents in the spinal CSF (typically lumbar injections), and tracking of drug concentration profiles in the spinal CSF, spinal tissue, cranial CSF, cranial tissue, blood and peripheral respectively^36^. Note that Eq (9) for the spinal CSF is an expanded version of simple diffusion eq (2) into which reaction and mass transfer terms have been incorporated.

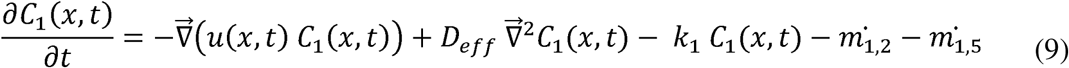

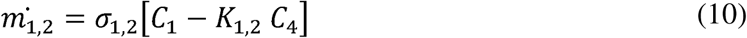

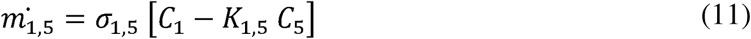

where *c*_1_(*x, t*) is the local concentration in the spinal CSF, *u*(*x, t*) is the local velocity which comes from injection, *D_eff_* is the *effective* diffusivity of the tracer in the CSF which is taken to be the same as *D* obtained experimentally via tracer distribution (cf Eq. 2). *k*_1_ is the first order reaction kinetics in the spinal CSF, *m*_1,2_ is the rate of absorption in the spinal tissue, and *m*_1,5_ is the rate of Antisense Oligonucleotides (ASO) leakage into the blood plasma. K is the distribution coefficient, a- is the mass transfer coefficient.

**Fig. 7.**
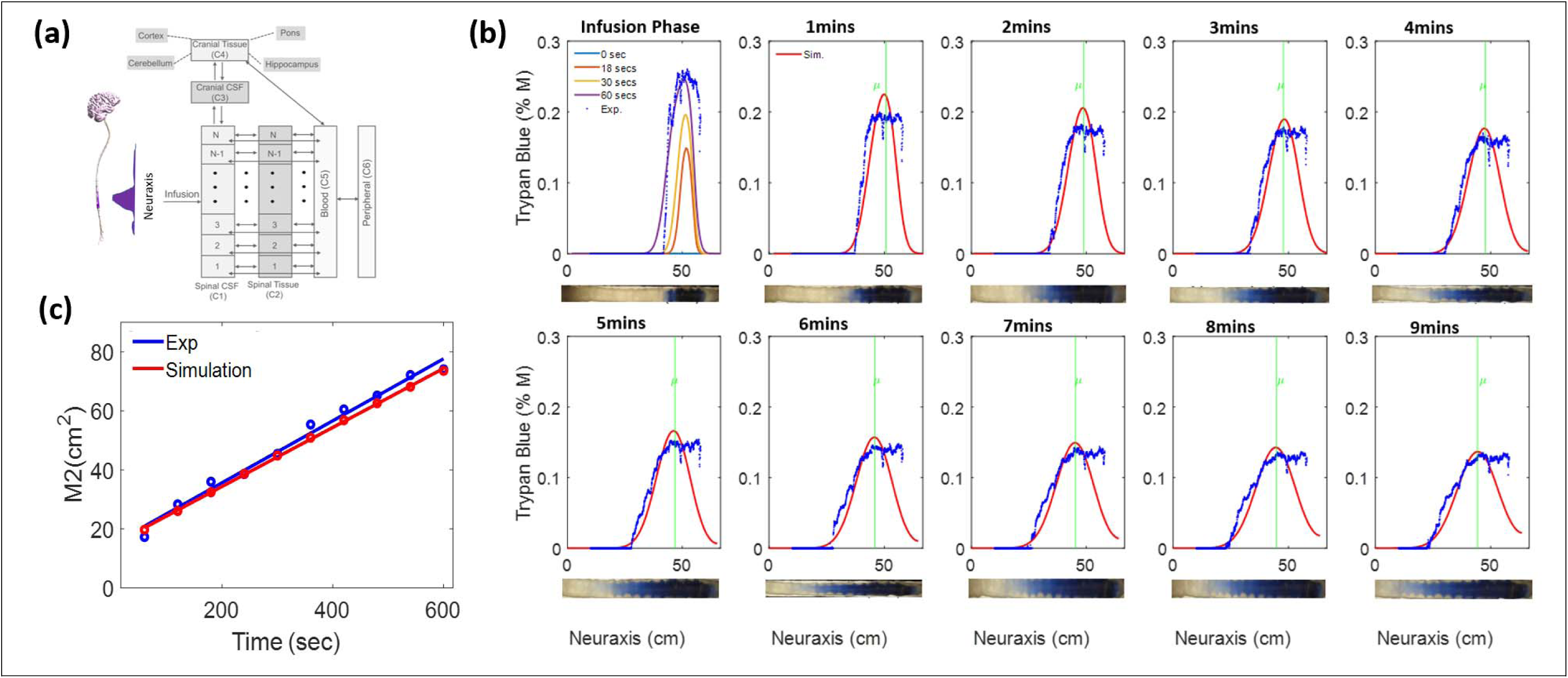
Figure showing the (a) schematic diagram used to derive the full pharmacokinetic model of drug dispersion. (b) evolution of the concentration profiles obtained using the pharmacokinetic model in Eq. (9) without mass transfer and reaction. (c) comparison of the second moment plots obtained from the experiment and simulation using the pharmacokinetic model in Eq. (9).

The PK model integrates the respective influences of physical drug transport (dispersion), biochemical reactions and mass transfer across CNS compartments (uptake). The left-hand side has drug accumulation (mol/min) expressed in terms of the drug concentration in the spinal CSF spaces, *c*_1_ (*x, t*). The convection flux is given by −*u*(*x, t*)*∇c1*(*x, t*), accounts for influences of high-volume injections on the CSF bulk flow velocity u(x, t) as well as effects of deformations during high volume injections. Drug dosing occurs via source terms at a specific location, x, and at instances in time according to the intrathecal injection source u(x,t). It varies in time at different locations, x, along the neuroaxis. CSF mediated *physical transport* is modeled as effective diffusive flux is given by ∇∇*D_eff_V c_1_*((*x, t*)], in terms of effective diffusivity, *D_eff_*, as determined in section 4.3.3. *Chemical kinetics* accounts for the conversion of active drug species. The reaction terms can also be used to account for enzymatic inhibition or drug binding on proteins which render the active agent inactive or immobile. *Mass transfer* fluxes measure the amount of drug that is eliminated from the CSF and transferred into spinal cord tissue (compartment 2) as shown in Fig. 7a. The mass transfer from CSF to spinal tissue is given in eq. (10). In addition, drug can be uptaken in epidural space, which can be considered a drug loss for the purpose of ASO therapy as in eq. (11). This PK model can account for clearance and biological activities. The PK model was used to simulate tracer profiles in our IT infusion experiments with reaction and mass transfer terms switched off, effectively replicating our bench tests in silico. Fig. 7b show the comparison between the profiles obtained experimentally and with those predicted with the pharmacokinetic model. Simulated concentration profiles track the optical intensity plots; second moment trajectories are in good agreement as seen in Fig 7c.

Simulating the pharmacokinetics of an ASO lead molecule over a period of several months of IT drug administration with multiple doses was possible in merely a few minutes with the proposed modeling framework^36^. Fig J1 in the supplement shows results demonstrating the ability to implement multiple feeds as well as data on how drug binding affects first and second moments. Unfortunately, detailed analysis of the interaction of transport and chemical kinetics is beyond the scope of this paper.

## 5 Discussion

Human CNS phantom. Using cast-and-mold techniques, we created a human spine model that is both anatomically and functionally accurate, including natural cerebrospinal fluid (CSF) pulsations with graded amplitudes. The phantom matches the real spine geometry as given by subject-specific MRI scans and reproduces tiny anatomical details like nerve roots. A key innovation of the CNS analogue is that it is fully sealed and deformable, which is crucial for experiments aiming to replicate oscillatory CSF motion with an amplitude that is graded along the entire neuroaxis of a dynamically deformable dural sack. Because the human spinal dura is flexible^42,43^, we also made the phantom’s dural boundary deformable, using a casting material with suitably configured mechanical properties. The casting material (TAP Platinum Silicone with tensile strength of 218 psi) makes the phantom tear-resistant and durable enabling it to withstand the stresses exerted by the pulsatile CSF motion, which is induced by forceful piston action of the pump. The transparent casting material does not interact with or entrain microscopic tracer particles; its adsorption inertia makes it dependable for optical particle tracking. By carefully 3D-printing of cast and molding with optimized catalyst mix ratio, the phantom achieves a precisely controlled thickness for the dura, excellent dimensional stability, with only 0.01% shrinkage. Controlling the thickness of the dural compartment in the manufactured phantom is essential, because its transparency and resistance to rupture both depend directly on it.

Applications. The proposed CNS replica mimics in vivo CSF dynamics and physical biodistribution in the closed and deformable spinal subarachnoid space in humans. The design permits realistic implementation and quantitative performance testing of infusion protocols used in the new drug trials^36^ or existing clinical practice for pain or spasticity management^32,44^. While this study focused on acute, high volume lumbar injection experiments aiming at maximizing caudo-cranial spread suitable for targeting the brain, the functional phantom can be deployed for bench trialing of novel multi-dosing regimes with high volume injections or chronic infusion protocols with drug pumps for individualized pain management. Thus, the human CNS phantom enables realistic performance testing and optimization of novel protocols, offering two key applications:

i. *Systematic study of effective biodispersion.* By adjusting CSF amplitude and frequency, we can systematically determine *effective dispersion* under realistic conditions of graded velocity profiles that are in effect globally across the neuraxis. This approach is a significant milestone toward validating predictions of theoretical models for drug dispersion in oscillatory flows^45^.
ii. *Design of new IT infusion protocols*. The transparent CNS surrogate allows for effective and inexpensive performance testing and optimization of new IT infusion protocols. The ability to manipulate infusion and physiological parameters easily, along with high-speed optical tracking of tracers, may in some cases be able replace animal trials, which do not scale to human anatomy. The CNS phantom is suited for conducting pilot experiments to develop and optimize infusion protocols before clinical trials. Among different injection techniques, high-volume injections are well suited for brain-targeted delivery because of the broader dispersion of the active pharmacological substrate. For a given injection time, high-volume infusions achieve larger dispersion distances than a drug pump, whose slow infusion rates would be relevant to spinal-located treatments.

Experimental determination of *effective* dispersion. The proposed bench setup enables adjustment of CSF flow dynamics—both amplitude and frequency—across a broad physiological range. Being able to freely vary CSF oscillations is critical for studying how changes in amplitude and frequency affect dispersion speed, which in turn allows describing physical transport using simple evidence-based formulae. The raw (dimensional) expressions for the dispersion coefficient effectively parameterizes the speed of biodispersion, making it ideal for clinicians who wish to estimate how fast substances spread in vivo. Additionally, two alternative formulations—derived through scale analysis—were introduced to express the dependence of observed tracer dispersion on dimensionless characteristics. The set of formulae can then be applied to predict *effective dispersion* during human intrathecal (IT) administration along the neuraxis with very little computational cost.

Image processing for tracer tracking tracer concentrations. The proposed image analysis protocol overcomes unavoidable noise in tracer intensity curves due to uneven surface or lighting conditions, and experimental variability. We previously had shown the beneficial use of tracking second moments^21^ to suppress errors in raw intensity curves. The slope of the temporal evolution of the second moment (M2) directly yielded estimates of the effective dispersion coefficient, *D*. However, due to asymmetry and boundedness of the confined spinal subarachnoid space, M2 trajectories depart from an ideal straight line as the tracer front progresses thus potentially introducing a bias in slope determination, which, in effect, impairs the experimental determination of *D*. The proposed new analysis based on moment inversion (MIP) eliminated this shortcoming. MIP overcomes the deficiency of the MoM helping to obtain better estimated of the true effective dispersion coefficient of the system. A comparison of the MIP to MoM results is given in Appendix G showing a deviation of up to 56.2%.

Pharmacokinetics. We also integrated biochemical interactions and drug uptake with physical transport by developing a comprehensive pharmacokinetic (PK) model for drug dispersion following IT administration. The PK model briefly introduced for the central CSF spaces in Eq. (9) enables analysis of drug distribution across various biological compartments and allows computationally inexpensive simulation of diverse dosing scenarios—including multiple dosing, CSF production, and flushing with artificial CSF to accelerate cranial delivery. Notably, PK simulations of IT administration of antisense oligonucleotides (ASO) for a period several months was possible in less than 10 CPU minutes. The PK model analysis also confirms that for rapid delivery to the brain, flushing with aCSF can be effective.

Nerve roots. We also confirmed experimentally that the effect of nerves roots is substantial. In a model without nerve roots, dispersion was four times slower than our main model with nerve roots. The experimental findings suggest that simplifying assumption of smooth annular cross-sectional profiles on which several theories are based do not capture significant geometry-induced mixing effects that occur only in the presence of microanatomical features. As a limiting point, our model does not contain dendrite ligaments and trabeculae, existing microfeatures in the spinal anatomy whose effects were addressed in a previous with in silico simulations^19^, showing to give rise to a 2.5 fold increase in local flow velocities. We chose not to include trabeculae and dendrite ligaments in our phantoms, because their number orientation and dimensions could not be resolved in our MR acquisitions.

Is IT dispersion a diffusive or convective process? Transport of solute in the human spinal canal has been studied by various authors^36,45,46^. Recently, several groups proposed theoretical models of drug dispersion in smooth idealized annular geometry without considering spinal microanatomy^47,48^. An elegant perturbation approach rendered time series results for species transport in the oscillatory fluid flow derived from the Navier Stokes equations. The net effect of CSF oscillations on species transport was purported as the combined effect of a Stokes drift^46^ and a steady streaming^45^ introducing *effective convective terms* into the reduced order species transport equations. The ability to match simulated concentration profiles obtained by direct numerical simulation (DNS) with results obtained with computationally much more efficient perturbation solutions (accounting for Stokes drift and steady streaming) significantly advance our theoretical understanding of species transport phenomena in oscillatory flow fields.

While the theory calls attention to *effective convection*, tracer dispersion observed in a complete CNS replica were consistently and adequately characterized by an *effective diffusion* process. We had previously stressed the significance of *geometry-induced mixing* around nerve roots, based on a series of experiments that center on the formation of eddies that form around microanatomical features^19,21,41^. Our experimental evidence suggests that pulsating flow around obstacles causes local micro-mixing which would have the globally observable effect of *diffusive (dispersive)* distribution of tracers molecules. It would be a significant development to characterize the global impact of microanatomy-like ligaments, nerve roots and trabeculae in theoretical approaches^47^. An important step in this direction is a recent work^50^, which appeared just after the completion of this study.

An outstanding question that remains to be shown is whether geometry-induced mixing can be incorporated into more complete theoretical descriptions. To facilitate the convergence of theory and CNS wide experiments, we generated parametric information of cross-sectional dimensions along the spinal neuraxis to enable future comparisons, but which are outside the scope of the current experimental study. The proposed elliptical shape approximations of the spinal anatomy might be useful to transfer data to theoretical models and validate simulation results against experimental evidence.

Our experiment also overcomes limitations of rigid models that require artificial manipulation of CSF pulsations or the lack features to study the effect of realistic infusion settings. Until open theoretical questions are settled and the effects of infusion mode and spinal deformations are incorporated into existing theoretical models, experimental evidence in combination with the proposed pharmacokinetic modeling framework of this study could serve as a guide for IT therapy design.

## 6 Conclusions

A functional anatomically accurate fully sealed and deformable model for human CNS was manufactured by a mold-and-cast technique. The manufactured phantom reproduces natural CSF pulsations with graded velocity profiles over a wide range of physiological conditions by emulating the expansion of the blood vasculature in the closed cranial compartment. Tracers were successfully injected and tracked optically within artificial CSF, to quantify the influence of infusion modes, drug dosage and subject-specific anatomy and pulsations. To the best of our knowledge, our model is the first closed deformable bench experiment capable of reproducing CSF dynamics driven by cranial blood/brain expansion within the entire human CNS. The functional CNS analogue might be useful for trial and to optimize infusion protocols in vitro before human trials and for generating drug dosing guidelines for IT administration using commercially available pumps and catheters.

## Acknowledge

We thank Dr. Charles Nicholson for fruitful discussions on image analysis of diffusive transport processes.

We also thank Brandon Donaldson, Kimberly Jacob-Paredes, and Gavin Enderlin for their contributions in image reconstruction.

